# patter: particle algorithms for animal tracking in R and Julia

**DOI:** 10.1101/2024.07.30.605733

**Authors:** Edward Lavender, Andreas Scheidegger, Carlo Albert, Stanisław W. Biber, Janine Illian, James Thorburn, Sophie Smout, Helen Moor

## Abstract

1. In the field of movement ecology, state-space models have emerged as a powerful modelling framework that represents individual movements and the processes that connect movements to observations. However, fitting state-space models to animal tracking data is often difficult and computationally expensive.
2. Here, we introduce `patter`, a package that provides particle filtering and smoothing algorithms that fit Bayesian state-space models to tracking data, with a focus on data from aquatic animals in autonomous receiver arrays. `patter` is written in `R`, with a high-performance `Julia` backend. Package functionality supports data simulation, preparation, filtering, smoothing and mapping.
3. In two worked examples, we demonstrate how to implement `patter` to reconstruct the movements of a tagged animal in an acoustic telemetry system from acoustic detections and ancillary observations. With perfect information, the particle filter reconstructs the true (unobserved) movement path (Example One). More generally, particle-based methods represent an individual’s possible location probabilistically as a weighted series of samples (‘particles’). In our illustration, we resolve an individual’s (unobserved) location every two minutes during one month in minutes and use particles to visualise movements, map space use and quantify residency (Example Two).
4. `patter` facilitates robust, flexible and efficient analyses of animal tracking data. The methods are widely applicable and enable refined analyses of home ranges, residency and habitat preferences.

## 1. INTRODUCTION

The field of movement ecology has attracted huge interest in recent decades (Nathan et al., 2008; Rafiq et al., 2021). An ever-increasing suite of electronic tagging and tracking technologies is used to track animals across the globe, providing a ‘panoramic window’ into their lives (Hussey et al., 2015). In aquatic environments, satellite tracking has reconstructed the migrations of air-breathing animals (Hays & Hawkes, 2018), archival geolocation has revealed the transoceanic movements of pelagic fish (Block et al., 2005) and autonomous receiver networks, known as passive acoustic telemetry arrays, have been established to track acoustically tagged benthic, demersal and pelagic taxa from local to continental scales (Matley et al., 2022). However, it is widely suggested that the accumulation of animal tracking data is outpacing the development of modelling methods and software packages for analysis (Rafiq et al., 2021).

In the last two decades, state-space models (SSMs) have emerged as a powerful modelling framework for animal tracking data (Patterson et al., 2008). An SSM is a hierarchical representation of a process-observer system in which the evolution of an unobserved (‘latent’) state (***s***) through time (*t* ∈ {1, …, *T*}) is imperfectly observed, generating ‘noisy’ observations (***y***_*t*_). In the context of animal tracking, SSMs model the movement process by which an animal’s location (***s***) evolves in (discrete) time (that is, *f*(***s***_*t*_ | ***s***_*t*−1_)) and the observation processes that connect movements to observations (that is, *f*(***y***_*t*_ | ***s***_*t*_)). The SSM thus forms a formal statistical framework within which it is possible to estimate the unobserved locations of an animal that are of interest, whilst accounting for properties of movement (including speeds and barriers) and observation processes (such as detectability). However, fitting SSMs can be challenging and computationally expensive (Patterson et al., 2008).

Particle filters are flexible Monte Carlo algorithms used to fit state-space models (Doucet & Johansen, 2009). In the context of animal tracking, a particle filter approximates the distribution of possible locations of an individual with a set of weighted samples termed ‘particles’ (Lavender et al., in prep). A movement model simulates possible locations of the individual (that is, ***s***_*t*_ ∼ *f*(***s***_*t*_ | ***s***_*t*−1_)) and observation model(s) weight particles in line with the probability of the observations (that is, *f*(***y***_*t*_ | ***s***_*t*_)). By resampling particles in line with the weights, we duplicate particles that are compatible with the data and eliminate incompatible particles. The result is an approximation of the distribution of individual’s location at each time step, given the preceding data (i.e., the partial marginal distribution, *f*(***s***_*t*_ | ***y***_1:*t*_)). Particle smoothers and samplers are extensions that approximate the full marginal (*f*(***s***_*t*_ | ***y***_1:*T*_)) and the joint (*f*(***s***_1:*T*_ | ***y***_1:*T*_)), respectively (Doucet & Johansen, 2009). Compared to alternative SSM-fitting methods for animal-tracking data, advantages of particle-based methods include their flexibility, scalability and the ease with which they can be intuitively understood. In the ecological literature, a handful of particle filtering routines have been developed, including for demersal fish geolocation at coarse spatial scales (over hundreds of kilometres) (Liu et al., 2019). However, existing routines are relatively specialised, computationally intensive and require significant user expertise.

Here, we introduce `patter`, an package that provides advanced particle filtering and smoothing algorithms for animal tracking data, motivated by our research in acoustic telemetry systems (Lavender, 2024a) (Lavender et al., in prep). The package is written in R (R Core Team, 2023) and integrates our high-performance `Julia` backend, `Patter.jl` (Lavender, 2024b). `Julia` is a relatively new language that combines the ease-of-use of an interpreted language like `R` with the speed of a compiled language like `C++` (Bezanson et al., 2017). `patter` includes routines for simulation, data preparation, particle filtering, smoothing and mapping. The package stands alongside generic SSM packages (King et al., 2016) but is designed for animal tracking—extending the `R` package ecosystem of this field (Joo et al., 2020) and that of the passive acoustic telemetry sub-field in particular (Kraft et al., 2023), with generic, flexible and fast particle algorithms. In the context of passive acoustic telemetry, `patter` is unique in the provision of routines that reconstruct individual movements and patterns of space use within a coherent probabilistic framework. The routines enable refined analyses of residency, home ranges and habitat preferences.

## 2. METHODOLOGY

### 2.1. Model formulation

The statistical methodology is described in Lavender et al. (in prep). This section provides a summary.

#### Posterior

We consider a Bayesian state-space model for the location of a tagged animal; that is, the joint distribution *f*(***s***_1:*T*_ | ***y***_1:*T*_), where in the simplest case ***s***_*t*_ = (*s*_*x*_, *s*_*y*_) is a two-dimensional vector that denotes location (in continuous, two-dimensional space), ***y***_*t*_ is the set of observations and *t* ∈ {1, 2, …, *T*} indexes time steps. The joint distribution is proportional to the product of a prior (the movement process) and the likelihood (the observation process), i.e.,

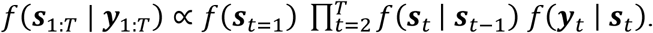

#### Prior

The prior comprises a probability density distribution of the animal’s initial location (*f*(***s***_*t*=1_)) and a movement model (*f*(***s***_*t*_ | ***s***_*t*−1_)). A simple model for *f*(***s***_*t*_ | ***s***_*t*−1_) in the two-dimensional case is an unrestricted, discrete-time random walk, where ***s***_*t*_ = (*s*_*x,t*−1_ +*d*_*t*_ cos *ϕ*_*t*_, *s*_*y,t*−1_ +*d*_*t*_ sin*ϕ*_*t*_) and *d* (step length) and *ϕ* (turning angle) are independently distributed random variables.

#### Likelihood

The likelihood measures the probability of the observations given the latent locations (***s***_*t*_). In an acoustic telemetry system, observations include acoustic measurements at each of *M* receivers (an *M* × *T* matrix, ***y***^(*A*)^), and/or ancillary data, such as depth measurements (a row vector, ***y***^(*D*)^). By way of example, here we consider a combined dataset ***y*** = {***y***^(*A*)^, ***y***^(*D*)^} and the likelihood 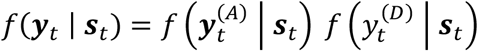

#### Acoustic measurements

The likelihood of the acoustic measurements at time 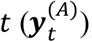, which comprise detections (1) or non-detections (0) at each operational receiver (that is, 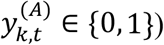 is modelled using the Bernoulli probability mass function, 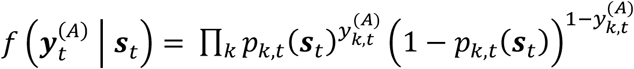 (assuming independence). We typically model detection probability, *p*, as a function of the distance between the receiver (at location ***r***_*k*_) and transmitter (at ***s***), such as, 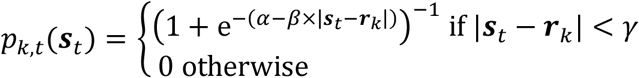 where α and *β* are parameters and *γ* is the detection range.

#### Depth measurements

A simple model for the likelihood of a depth observation is:

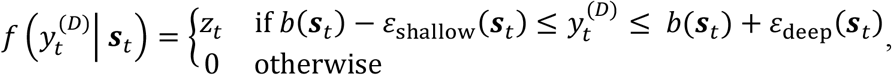

where *z*_*t*_ = (*ε*_deep_ (***s***_*t*_) +*ε*_shallow_(***s***_*t*_))^−1^. This requires 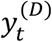 to be within an envelope around the bathymetric depth *b*(***s***_*i*_) defined by shallow- and deep depth-adjustment functions, *ε*_shallow_(***s***_*t*_), *ε*_deep_(***s***_*t*_) ≤ *b*(***s***_*t*_). These functions represent observational and spatially explicit bathymetric uncertainty and can be tailored for different species. For benthic species, small errors (*ε*_shallow_(***s***_*t*_), *ε*_deep_(***s***_*t*_) ≪ *b*(***s***_*t*_)) only permit observations close to the seabed; for pelagic species larger *ε*_shallow_(***s***_*t*_) values permit observations in the water column.

### 2.2. Model inference

#### Filter

For inference, we begin with the partial marginal distribution, *f*(***s***_*t*_ | ***y***_1:*t*_). Particle filters approximate *f*(***s***_*t*_ | ***y***_1:*t*_) as a sum of *N* weighted particles, i.e., 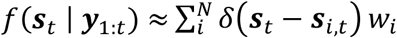, where *δ* is the Dirac delta function, *w* denotes normalised weights and *i* indexes particles. We assume static parameters are known. Starting with an initial set of particles sampled from the prior (i.e., ***s***_*i,t*=1_ ∼ *f*(***s***_*t*=1_)), the filter iteratively simulates particles via the movement model (i.e., ***s***_*i,t*_ ∼ *f*(***s***_*i,t*_ | ***s***_*i,t*−1_)), weights particles in line with the likelihood (via *w*_*i,t*_ ∝ *w*_*i,t*−1_*f*(***y***_*t*_ | ***s***_*i,t*_)) and resamples particles accordingly.

#### Smoother

Particle smoothers re-weight filtered particles to approximate the full marginal, *f*(***s***_*t*_ | ***y***_1:*T*_). The two-filter smoother uses ***s***_*i*,…,*N,t*_ particles from a forward filter (with weights *w*_*i,t*_) and 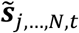 particles from a backward filter (with weights 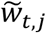). The distribution *f*(***s***_*t*_ | ***y***_1:*T*_) is approximated as a sum of re-weighted particles via 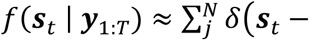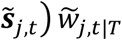 where the smoothed weights for each particle 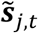 represent a summation over all possible movements from preceding particles on the forward filter, that is,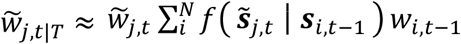. We use samples from *f*(***s***_*t*_ | ***y***_1:*T*_) to map utilisation distributions (Lavender et al., in prep). Sampling efficiently from the joint distribution, *f*(***s***_1:*T*_ | ***y***_1:*T*_), is more challenging and beyond the scope of this contribution.

## 3. PACKAGE

`patter` supports data input, algorithms and mapping (Fig. 1).

**Figure 1.**
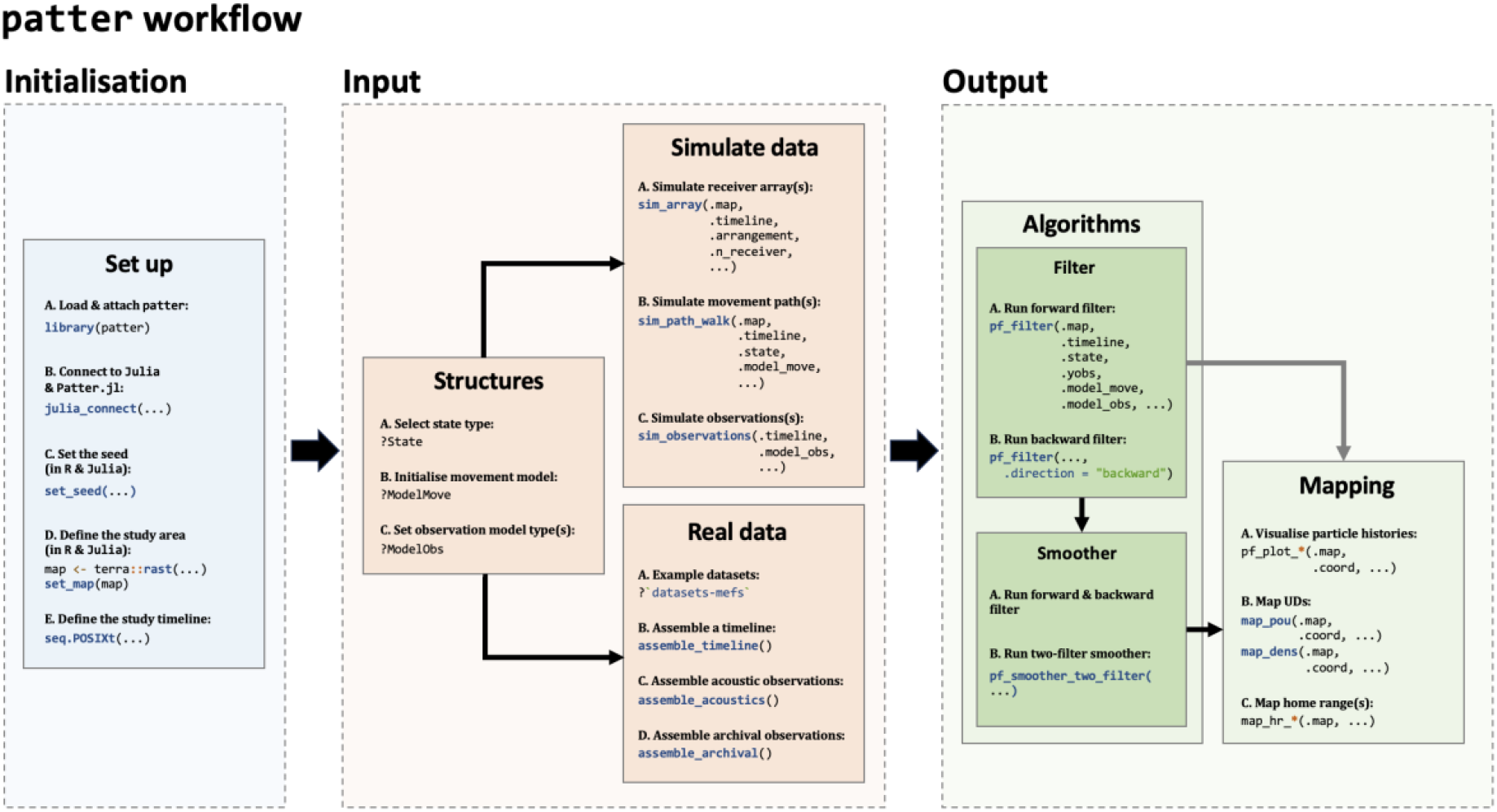
Overview of the `patter` package.

For data input, `patter` provides `sim_*()` functions for *de novo* simulation or accepts real-world datasets.

Particle algorithms are implemented by `pf_*()` functions. Filtering and smoothing are implemented via `pf_filter()` and `pf_smoother_two_filter()`. We provide movement models and methods that evaluate the likelihood of acoustic and archival (depth) observations, but algorithm components can be customised as required. The main output of particle routines is a `data.table` of particles. The core algorithm routines wrap our high-performance `Patter.jl` package. `JuliaCall` implements the coupling between `patter` and `Patter.jl` (Li, 2019). Movement and observation models are multithreaded and designed for numerical stability. We anticipate that most users will prefer the `R` front-end, but `Patter.jl` can also be used directly.

Mapping functions (`map_*()`) facilitate subsequent analysis, including the reconstruction of utilisation distributions.

## 4. EXAMPLES

### 4.1. Overview

We provide two worked examples using simulated data. Simulation is a useful starting point for illustration and real-world analyses, where it informs algorithm implementation and interpretation. In both examples, we consider the movements of a benthic animal in a hypothetical acoustic array spanning a Marine Protected Area (MPA) in Scotland (Fig. 2). The study area is defined by a 100 × 100 m bathymetry grid. We base the grid on real-world data (Howe et al., 2014) but for the purposes of our first example add some random noise such that each cell’s depth is unique. Within this region, we tag an animal with an acoustic transmitter and an archival (depth) tag. We imagine that both tags operate at a resolution of two minutes and simulate a discrete-time random walk at this resolution over a one-month period (Fig. 2B–C). We simulate acoustic and depth observations arising from the simulated path and apply our algorithms to these data to reconstruct movements and patterns of space use. In the first example, we simulate observations in such a way that the depth observation exactly defines the location of the individual (which is situated on the seafloor) (Fig. 2D–F). This example will demonstrate that in the absence of uncertainty the particle filter reconstructs the true movement path. In the second example, we simulate observations probabilistically and demonstrate the representation of uncertainty in particle-based methods and the reconstruction of maps of space use (Fig. 2D–F). In the following sections, we showcase key functions and arguments (Fig. 1). Additional arguments are denoted by ellipses. Complete code is available online (Lavender et al., 2024).

**Figure 2.**
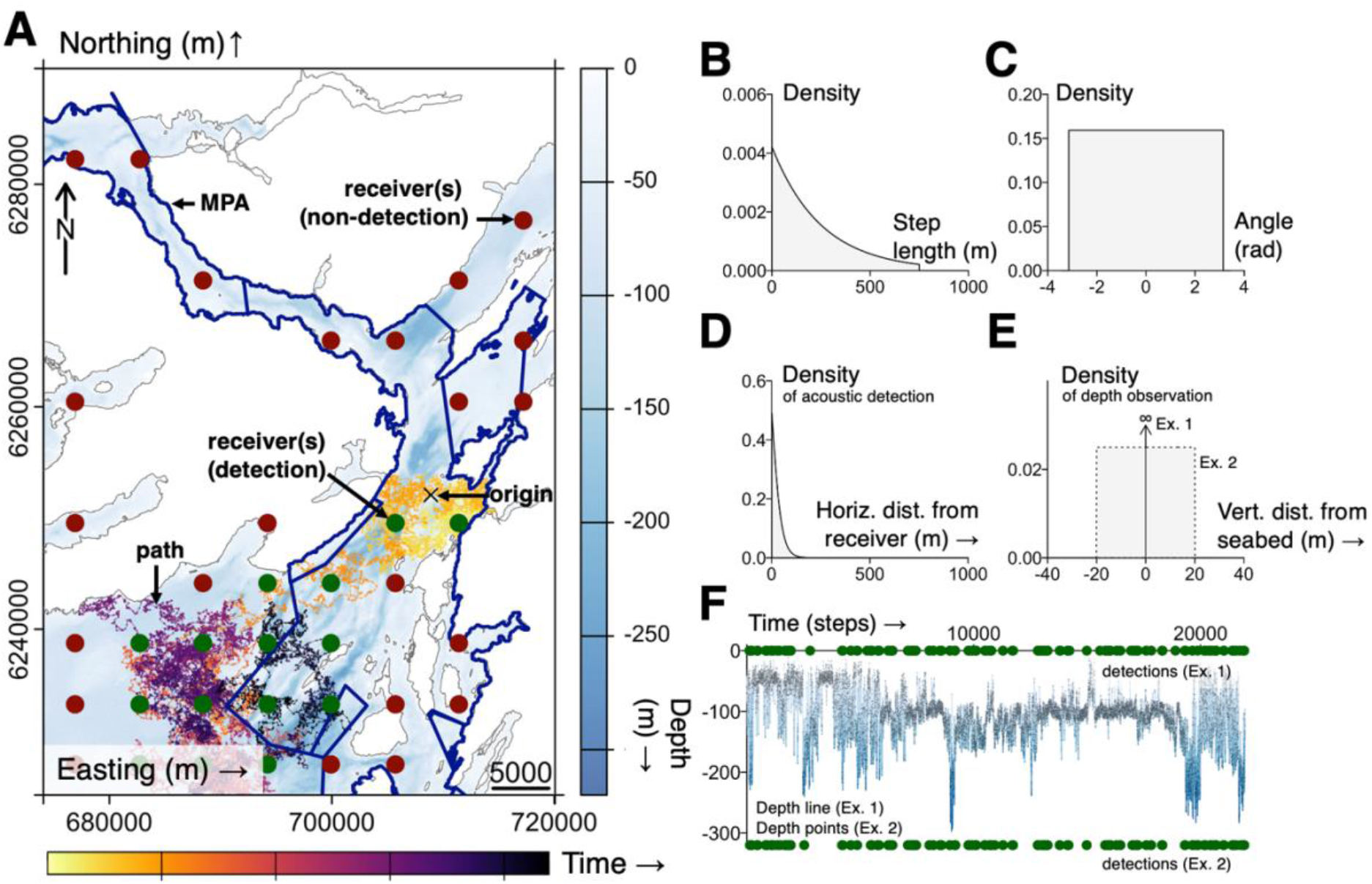
The components of a state-space model for animal tracking data. **A** shows the study area, including a simulated movement path (coloured by time) and acoustic receivers (sized by detection range and coloured by detection(s)/non-detection). **B–C** show the components of the random walk used to simulate and model movements. **D–E** show the observation models used to simulate and model acoustic and depth observations arising from the simulated path in the two worked examples; the acoustic observation model is constant but the depth model differs (see §4.1). **F** shows the simulated time series for each example.

### 4.2. Implementation

#### 4.2.1. Simulation

We begin with essential initiation:

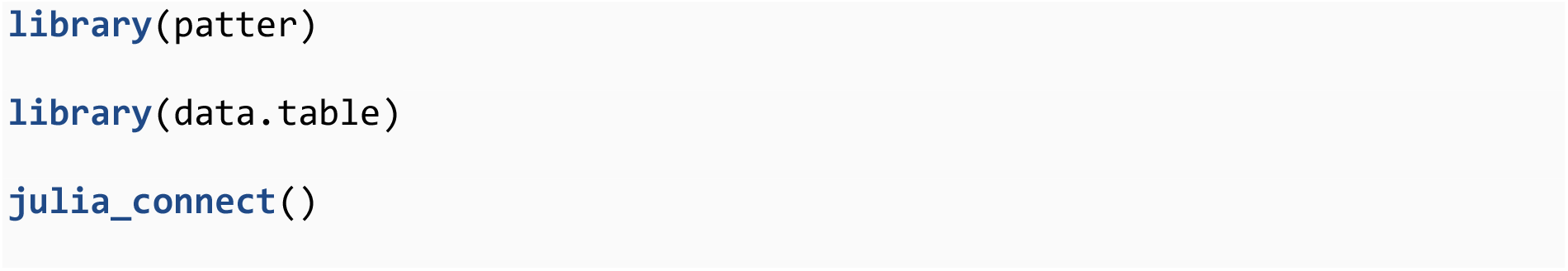

Next, we define the study system:

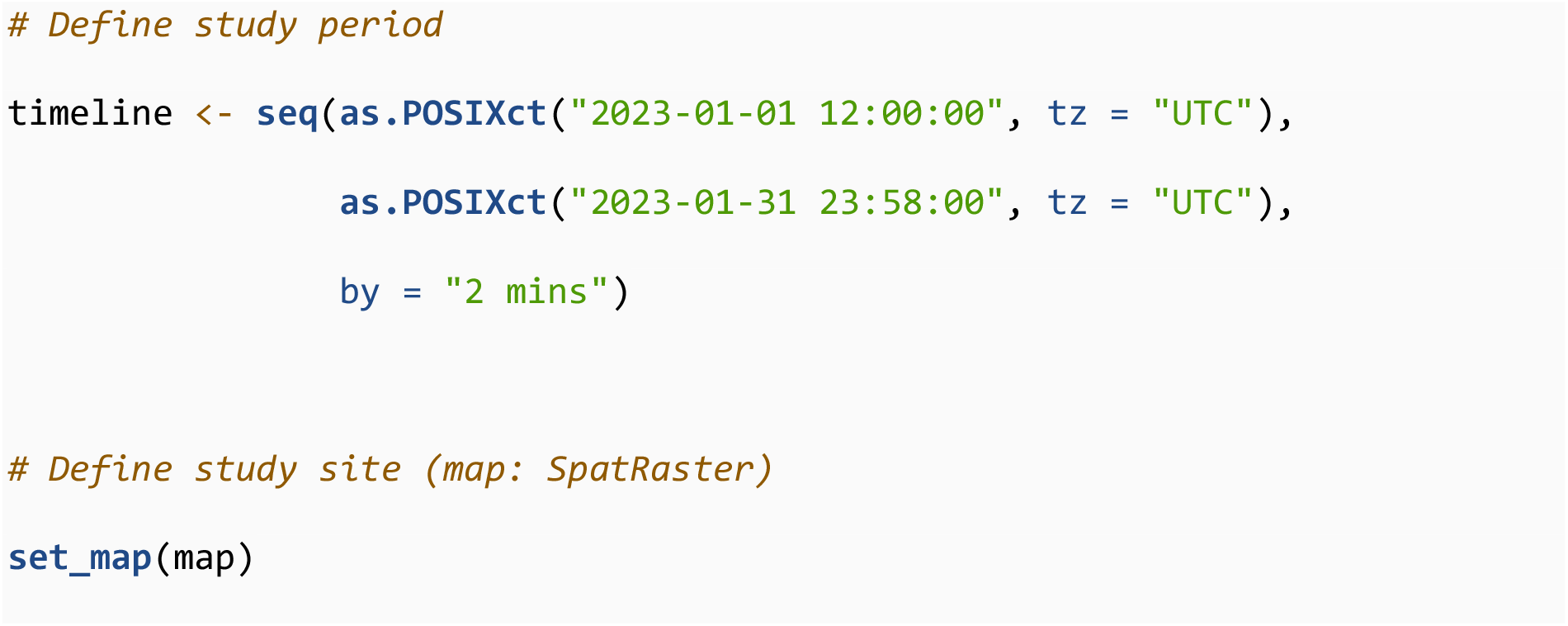

We simulate an acoustic array (i.e., `data.table` of receivers). We include three `receiver` columns that represent observation model (detection probability) parameters:

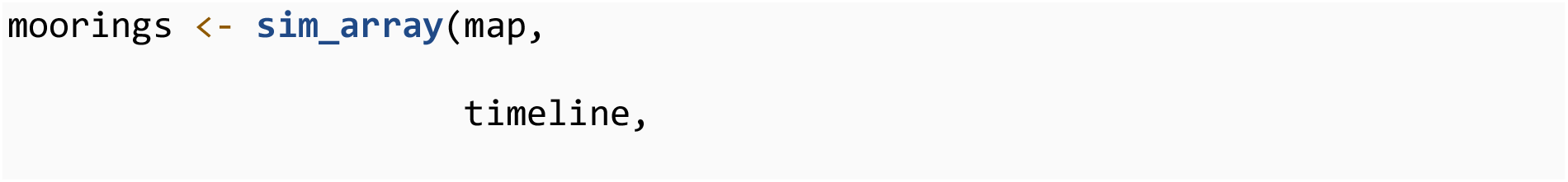

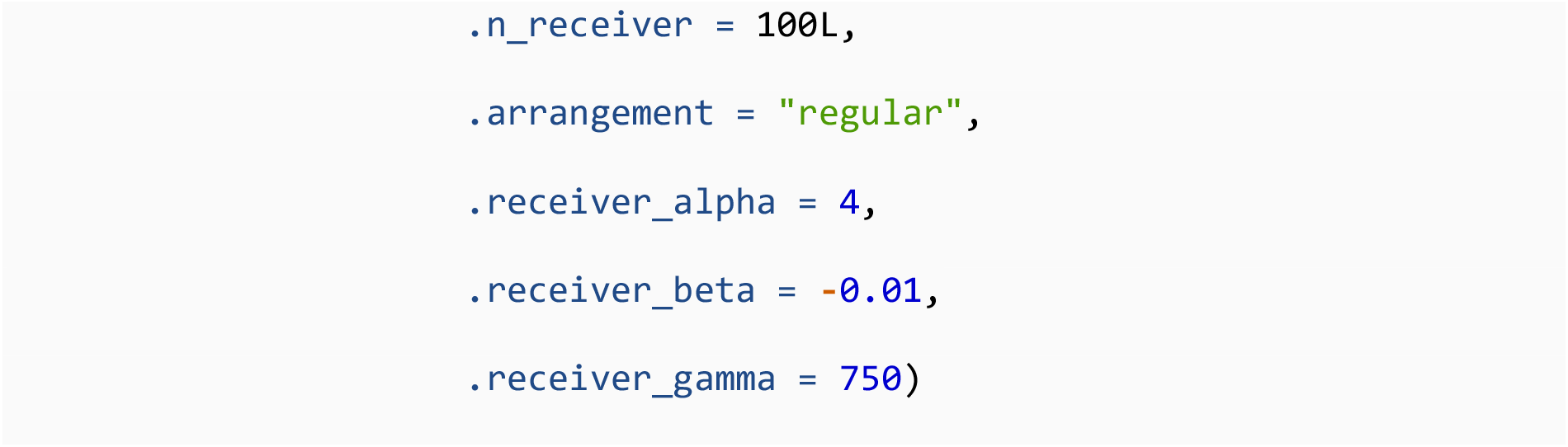

We then simulate a two-dimensional random walk in this area:

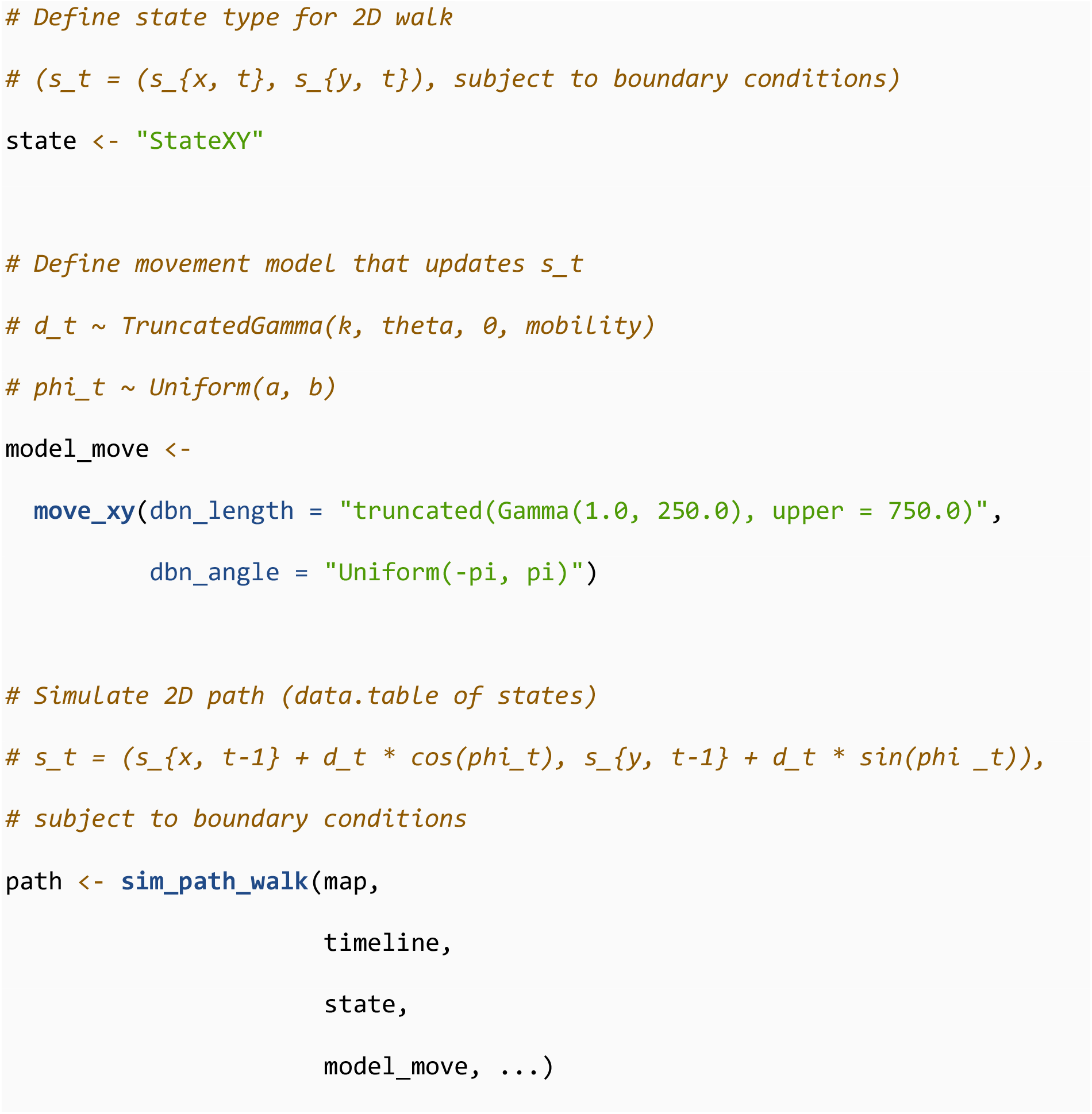

Next, we simulate acoustic and archival observations. This requires defining a vector of observation model (`ModelObs`) structures (which hold model parameters) and a corresponding `list` of `data.tables` (with those parameters). For both examples, we simulate acoustic observations from a truncated logistic model (for which we provide the `ModelObsAcousticLogisTrunc` structure and the essential parameters are defined in `moorings`). To simulate depths, we consider a simple version of the uniform model described previously, as implemented by the `ModelObsDepthUniform` structure, where *ε*_shallow_(***s***_*t*_) and *ε*_deep_(***s***_*t*_) are the constants depth_shallow_eps and dep*t*h_deep_eps.

In the first example, we imagine an animal found exclusively on the seabed, the depth of which is known exactly, giving parameters:

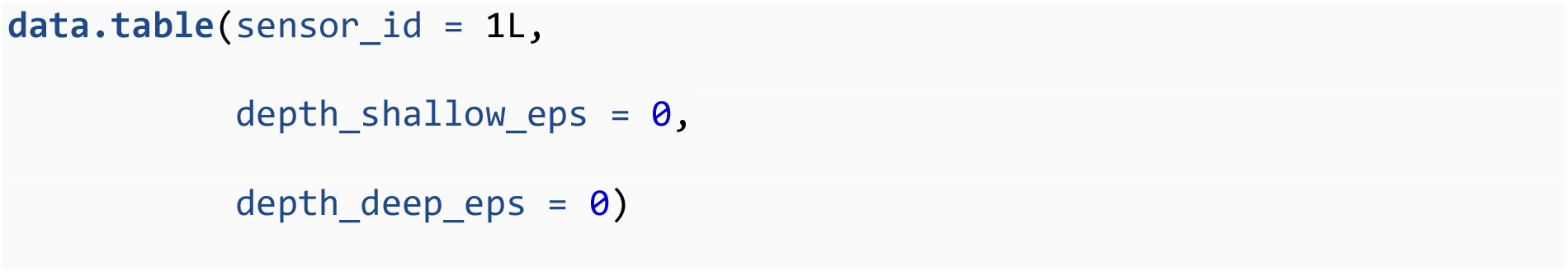

In the second example, we incorporate uncertainty:

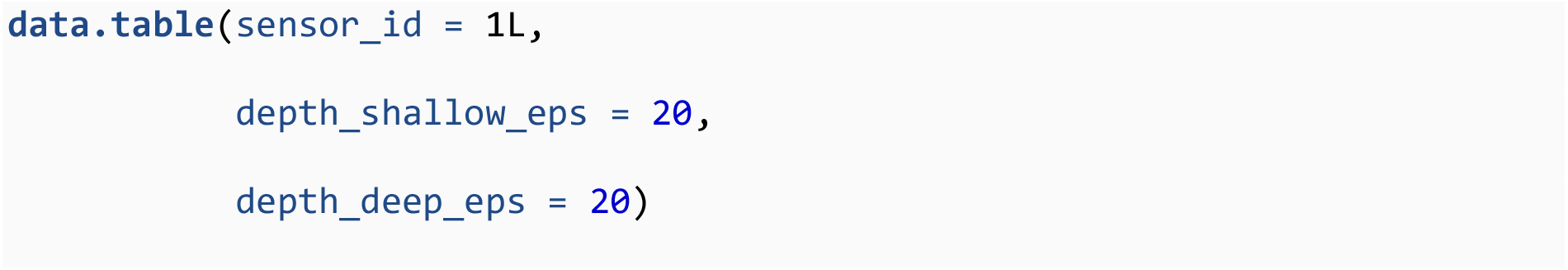

For each example, `sim_observations()` simulates a `list` of observations:

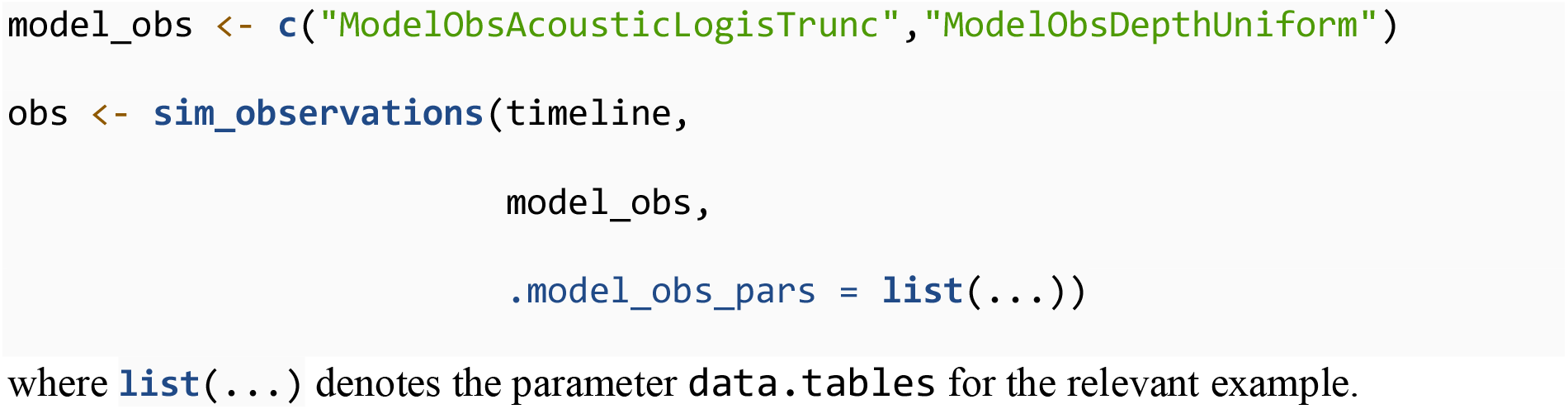

where **list**(…) denotes the parameter `data.tables` for the relevant example.

#### 4.2.2. Particle filter

The particle filter is implemented via:

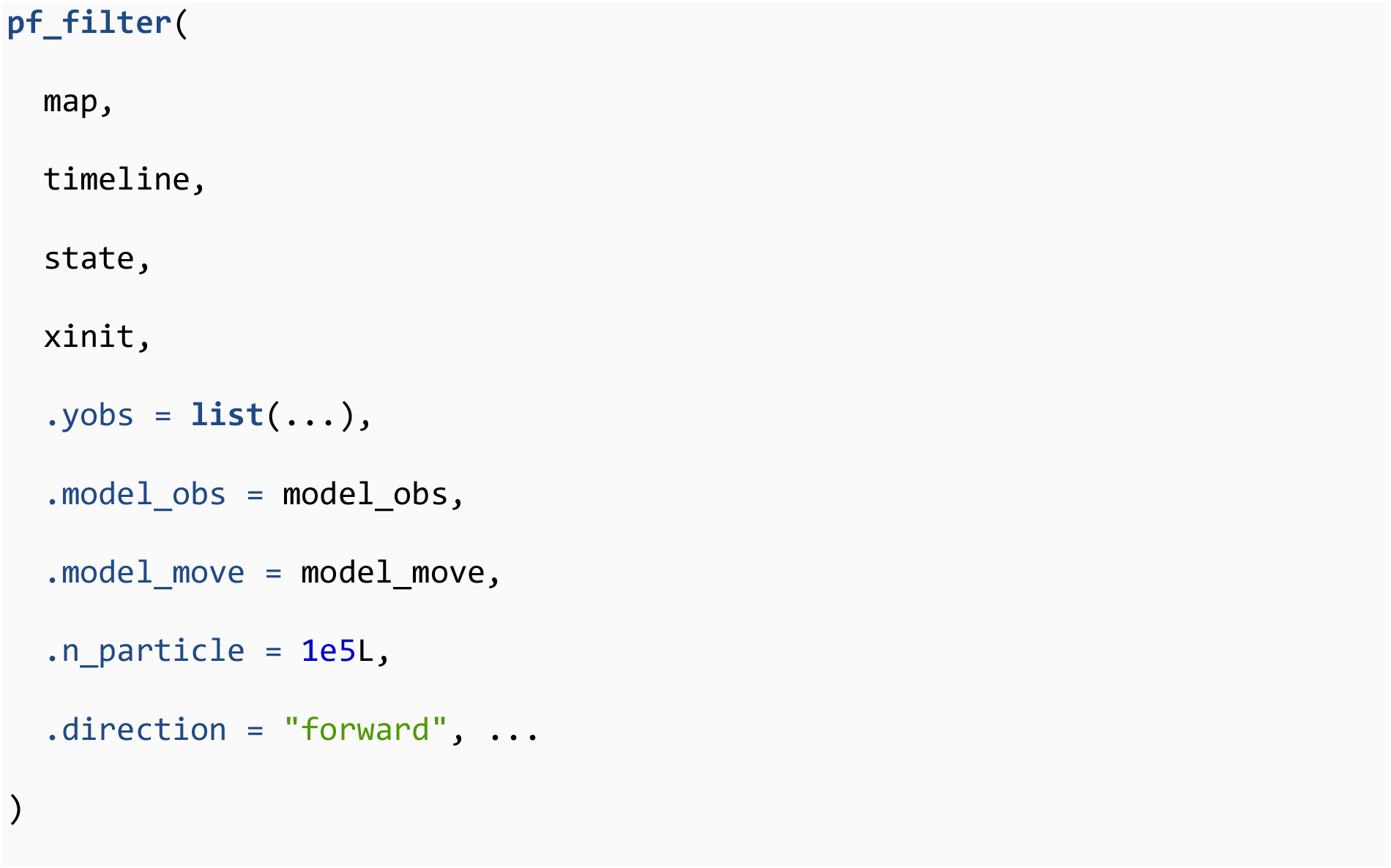

where `.xinit` (optional) is the simulated tagging location and `.yobs` is the `list` of datasets for the relevant example. This returns a `pf_particles`-class object that includes a `data.table` of particles and diagnostic statistics. In the first example, the filter reconstructs the true (unobserved) path. In the second example, we generate a ‘cloud’ of particles at each time step, for which we examine particle diagnostics and proceed to smoothing.

#### 4.2.3. Particle smoother

Particle smoothing is implemented using outputs from a forward and backward filter via:

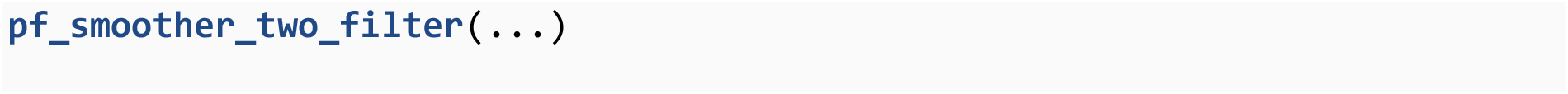

For illustration, we reconstruct the utilisation distribution and home range from smoothed particles via `map_dens()` and `map_hr()`. We also compare time spent in the MPA, estimated from the proportion of particles inside the MPA, to the truth.

This workflow is highly customisable. Users can define species-specific movement models (of any dimension), include diverse observation types and implement system-specific observation models. See the package documentation for details.

#### 4.3. Results

In our first example, the particle filter reconstructs the simulated movement path perfectly (Fig. 3A). In the second example, in which observations were simulated with error, the particle filter represents the individual’s possible locations at each time step with a series of weighted particles that approximate the partial marginal distribution, *f*(***s***_*t*_ | ***y***_1:*t*_) (Fig. 3B). The particle smoother re-weights filtered particles, approximating the full marginal, *f*(***s***_*t*_ | ***y***_1:*T*_) (Fig. 3C).

**Figure 3.**
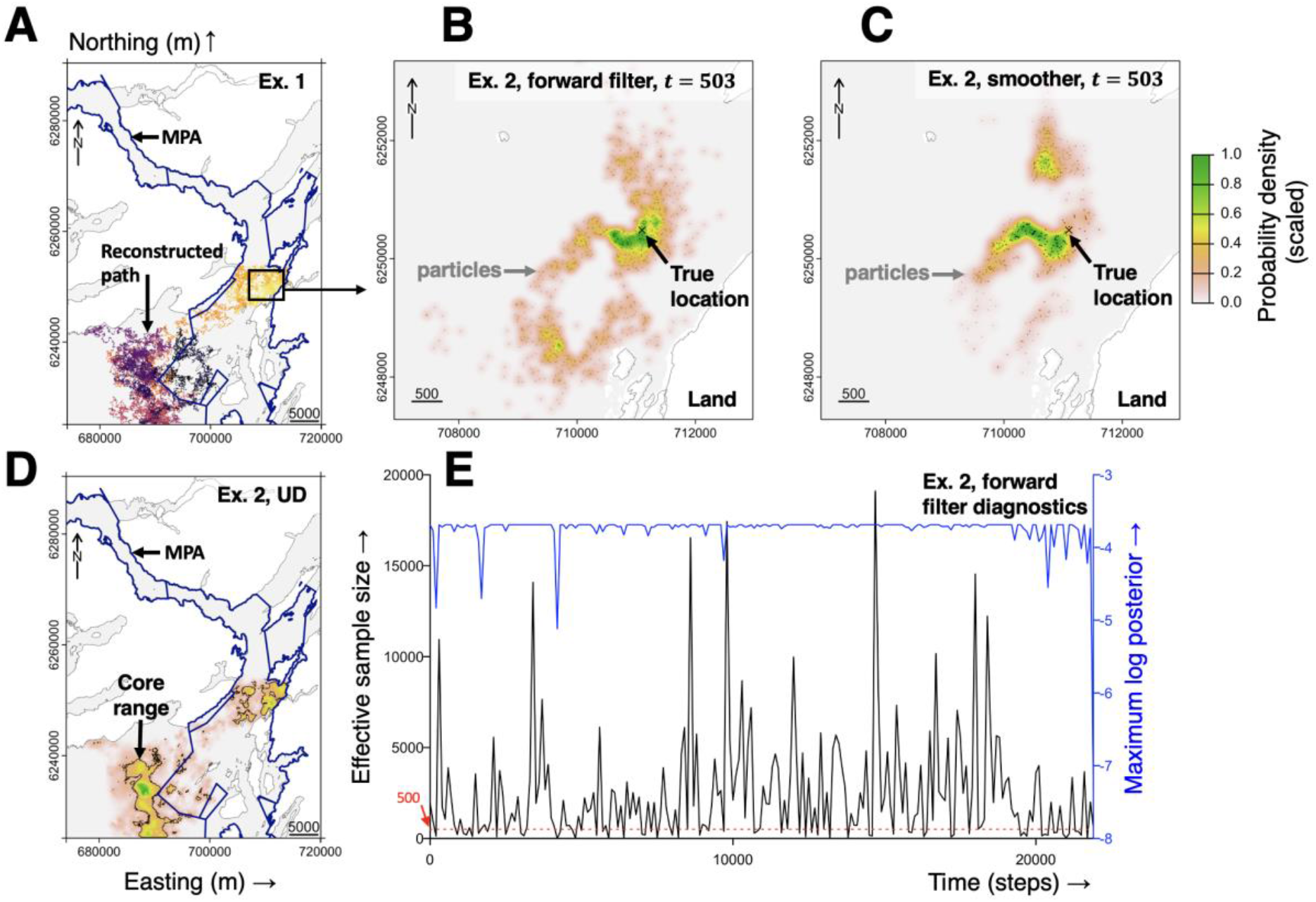
patter outputs for the first (A) and second (B–E) worked examples. **A** is the movement path reconstructed by the particle filter for the first example. The simulated path is recovered perfectly (at grid resolution) because the observations exactly define the individual’s location. **B–C** show particles and scaled probability densities from the forward filter and two-filter smoother, respectively, at a selected time step. **D** maps the pattern of space use over the entire time series using smoothed particles. Core ranges contain 50 % of the probability mass volume. Estimated residency in the MPA is 37.0 % (the true value is 36.3 %). **E** shows diagnostics from the forward filter. The minimum value of each statistic is shown for every 100 time steps.

Smoothed particles can be used to map patterns of space use, estimate home ranges and quantify residency (Fig. 3D). The quality of the smoothing depends on the filter. In this case, filter diagnostics are adequate (Fig. 3E). Total computation time ranged from 5–32 minutes for examples 1–2 on a 2023 MacBook Pro (Apple M2 Pro, 32 GB RAM, 12 CPUs).

## 5. DISCUSSION

`patter` provides a robust, fast and accessible implementation of particle-based methods for animal tracking data (Lavender et al. in prep). In the context of passive acoustic telemetry, `patter` is unique in the provision of algorithms that represent the movement and observation processes that generate observations, including movement capacity, barriers to movement, detections and ancillary observations, within a biologically and statistically sound framework. The algorithms outperform standard heuristic analytical approaches (provided by packages such as `VTrack` and `RSP`) (Lavender et al., in prep). Furthermore, the movement and observation models are customisable, which makes the routines applicable in many real-world settings. However, understanding the settings in which different methods are more or less useful remains an important research area (Lavender et al., in prep).

A key feature of `patter` is the speed provided by the `Patter.jl` backend (Lavender, 2024b). Previous research has linked the underutilisation of particle algorithms in ecology to their computational requirements (Liu et al., 2019), which are typically *𝒪*(*NT*) for filtering and *𝒪*(*N*^2^*T*) for smoothing (Doucet & Johansen, 2009). While adequate filtering is often possible with relatively few particles (*N* ≈ 1000), this imposes some practical constraints that can be limiting in situations where numerous particles are required to ensure convergence (namely, labyrinthine landscapes where a tiny fraction of possible routes are compatible with the data) (Lavender et al., in prep). The particle penalty is more severe for smoothing but mitigated via subsampling, since in general only a subset of the particles required for a successful filter run are ultimately required to approximate the distribution of latent locations (in two or three dimensions). In practice, our packages achieve speeds that compare favourably with other geolocation routines (even with ≤ 1 million particles). For example, Hostetter & Royle (2020) formulated a state-space model for acoustic detections (*T* = 150) alongside a bespoke Jags implementation that requires ∼15 hours on a standard computer to run. With our particle filtering–smoothing algorithm, the estimation of latent locations in this situation is soluble in seconds. Other geolocation packages developed for demersal fish that fit hidden Markov models using maximum likelihood typically require hours or days to derive daily geolocation estimates for a one-year period and even a novel Python particle filter that exploits GPU-parallelisation takes up to an hour (Liu et al., 2019). While run times are not directly comparable, it is encouraging to see `patter` achieving speeds sufficient to make particle algorithms serious candidates for real-world analyses (Lavender et al., in prep).

For applied studies, we suggest the use of simulations to guide method implementation and interpretation. Particle filters can be sensitive to model parameters and tuning settings (such as particle number) and system-specific simulations can inform input arguments and quantify sensitivity (Lavender et al., in prep). Joint estimation of model parameters and latent locations is a possible future development. As an example real-world analysis, we are currently analysing acoustic and archival data from the Critically Endangered flapper skate (*Dipturus intermedius*) to quantify patterns of space use and site affinity to a Scottish Marine Protected Area. There is much to be learnt from applications in other settings and we welcome community feedback as these developments are exploited.

## Acknowledgments

EL was supported by a postdoctoral researcher position at Eawag, funded by the Department of Systems Analysis, Integrated Assessment and Modelling.

